# Replication poison treated BRCA1-deficient breast cancers are prone to MRE11 over-resection resulting in single strand DNA accumulation and mitotic catastrophe

**DOI:** 10.1101/2023.10.31.564929

**Authors:** Imene Tabet, Esin Orhan, Ermes Candiello, Lise Fenou, Carolina Velazquez, Beatrice Orsetti, Geneviève Rodier, William Jacot, Cyril Ribeyre, Claude Sardet, Charles Theillet

## Abstract

BRCA1, BRCA2 and RAD51, key players of homologous recombination (HR) repair, are also involved in stalled DNA replication fork protection and repair. BRCA1-deficiency is encountered in 25% of Triple Negative Breast Cancer (TNBC). Here we investigated the sensitivity of *BRCA1*-deficient TNBC cell models to gemcitabine a frequently used replication poison that does not alter DNA structure. We show that BRCA1-deficient models, in contrast to their isogenic *BRCA1*-proficient counterparts, are superiorly sensitive to gemcitabine, accumulate massive levels of single strand DNA (ssDNA), in absence of RPA and RAD51 signals and elevated double strand break (DSB) numbers leading to cell death.

Remarkably, ssDNA accumulation in gemcitabine-treated BRCA1-deficient cells was strongly diminished by the MRE11 inhibitor mirin, while it did not affect ssDNA levels resulting from PARP inhibitor olaparib treatment. The central role of MRE11 DNA resection strongly suggested that replication fork reversal may be important in response to replication poisoning by gemcitabine in BRCA1-deficient models. Furthermore, we demonstrate that gemcitabine-treated *BRCA1*-deficient cells showing massive ssDNA accumulation slipped into mitosis and produced mitotic bridges and micronuclei (MN) showing strong BrdU and γH2AX staining. Noticeably these BrdU-positive MN and DNA bridges triggered cGAS sensing.

Our data, thus, strongly suggest that gemcitabine treatment could be beneficial in *BRCA1*-deficient TNBC both in terms of cancer cell death, but possibly as well in terms of antitumor immune response.

## Introduction

Homologous recombination deficient (HRD) cancers, originally identified in a subset of ovarian and breast cancers with pathological *BRCA1* or *BRCA2* mutations, are characterized by an elevated genetic instability and increased sensitivity to platinum salts and PARP inhibitors (PARPi) (1). The exquisite sensitivity to genotoxic treatment of BRCA-deficient tumors has long been considered to result from faulty repair of drug induced double strand DNA breaks (DSB). However, recent data suggested that central HR players such as *BRCA1*, *BRCA2* and *RAD51* could also play important roles in the protection, repair and restart of stalled DNA replication forks (2), (3). Moreover, BRCA-deficient models exhibit diminished fork progression speed and accumulate single strand DNA (ssDNA) at replication forks, even in absence of exogenous stress (4), (5). Altogether, this supports the idea that BRCA-deficient tumors could be at particular risk of undergoing DNA replication malfunction and, thus, be particularly sensitive to replication poison-based treatment. DNA replication fork stalling can result from multiple causes, but in case of exposure to anticancer drugs mainly derives from obstacles hindering fork progression or from nucleotide pool imbalance (6).

Depending on the number of stalled replication forks and repair capacity, extended replication stress can result in ssDNA accumulation and affect cell viability.

Triple Negative Breast Cancer (TNBC) represent 15% of all breast cancers and its most aggressive subtype. Noticeably, TNBC comprises up to 25% of *BRCA1-*deficient tumors due either to pathological coding mutations or epigenetic silencing of the promoter region (7), (8).

Based on the prevalence of *BRCA1* deficiency in TNBC, we, thus, undertook to study the impact of acute replication stress induced by the replication poison gemcitabine in *BRCA1*-deficient models, in comparison with their *BRCA1*-proficients counterparts. Gemcitabine is commonly administered in cancer care and, noticeably, in TNBC recurrence. Moreover, our interest in gemcitabine stemmed from the fact that, in contrast with other genotoxic drugs, it is not known to alter the DNA structure. Indeed, as an irreversible inhibitor of the large subunit of ribonucleoside-diphosphate reductase (RRM1), it impairs pyrimidine biosynthesis and produces nucleotide pool imbalance. Furthermore, being incorporated in nascent DNA, it acts as a chain terminator (9). We show here that, in comparison with their cognate *BRCA1*-proficient cells, *BRCA1*-deficient models are superiorly sensitive to gemcitabine, presenting a large fraction of cells that accumulate massive levels of single strand DNA (ssDNA), in absence of the ssDNA binding protein RPA signals. Ultimately these cells undergo elevated DSB numbers and die. Our data show that the massive accumulation of ssDNA in a *BRCA1*-deficient context is associated with uncontrolled DNA hyper-resection by the MRE11 exonuclease. MRE11 is instrumental in the onset of DNA HR repair, being actively recruited onto DNA DSB by NBS1/CtIP, where it produces the overhangs necessary for homologous recombination (10). Interestingly, MRE11 has also been shown to be recruited by NBS1/CtIP onto stalled and reversed replication forks where it resects nascent DNA (11), (12). Furthermore, we demonstrate that *BRCA1*-deficient cells, undergoing ssDNA accumulation, slip into mitosis, produce aberrant mitoses and micronuclei (MN) showing strong ssDNA-specific BrdU and γH2AX staining. Furthermore, these MN presented a strong cyclic GMP-AMP synthase (cGAS) signal, suggesting they could participate in the activation of innate immunity.

Overall, our data show that *BRCA1*-deficient cancer cells are prone to undergo lethal ssDNA accumulation when treated with gemcitabine and, thus, suggest that replication poisons could have a favorable therapeutic impact on these tumors possibly assorted to innate immunity activation.

## Materials and methods

### Cell lines and CRISPR-Cas9 engineered mutants

SUM159 TNBC cells were a gift from Dr S Ethier (MUSC, Charleston, SC), MDA-MB436, BT549, HCC38 OVCAR8, UWB1-289 and its variant UWB1-289B1 ectopically expressing a full length BRCA1 construct, were obtained from ATCC®. The CRISPR-Cas9 BRCA1-KO clone was engineered in house from SUM159PT. SUM-159 were maintained in Ham’s F-12 medium (Gibco™, Fisher Scientific, Illkirch-Graffenstaden, France) supplemented with 5% FBS, 10 µg/ml insulin, 1µg/ml hydrocortisone and 1% Antibiotic-antimycotic (100X) (Gibco™, Fisher Scientific, Illkirch-Graffenstaden, France), UWB1.289PT in the 50% RPMI-1640 (Gibco™)/ 50% MEGM (MEGM Bullet Kit; CC-3150, Lonza, Basel, Switzerland) supplemented with 3% FBS, 1% Antibiotic-antimycotic (100X) (Gibco™, Fisher Scientific, Illkirch-Graffenstaden, France), HCC38, BT-549 and OVCAR8 in RPMI-1640 supplemented with 10% FBS and 1% Antibiotic-antimycotic. All cell lines and selected clones were genetically typed by Eurofins Genomics cell line authentification (Eurofins Genomics, Les Ulis, France).

### Immunofluorescence

Cells were grown on 12mm diameter slides cover slips in 24 well-plate for 24h, then drugs (gemcitabine or alaparib) were added at predetermined IC50 concentrations (Lilly, Fegersheim, France). Cells and tumor sections were sequentially subjected to mild pre-extraction (PBS 0.4% Triton X100, 5min at 4°C), fixation (PBS 4% PFA) and blockage/permeabilization (PBS 3% BSA + 0.2% Triton X100, 1h at room temperature), incubated overnight at 4°C with the primary antibody (diluted in PBS 3% BSA + 0.2% Triton X100), then with the secondary antibody (diluted in PBS 3% BSA + 0.2% Triton X100, 1h at room temperature). Between each step, slides were washed 3 times with PBS. Cells were counterstained with DAPI (Fisher Scientific, Illkirch-Graffenstaden, France) and mounted with MWL4-88 cover slips (Citifluor, CliniSciences, Nanterre, France) and stored at 4°C. Antibodies are described in the Antibody section. Single strand DNA (ssDNA) detection experiments were performed on cells incubated with 10μM BrdU (Sigma Aldrich, Saint Quentin Fallavier, France) for 24h prior gemcitabine treatment. BrdU was revealed with an antibody in non-denaturing conditions. In MRE11 inhibition experiments cells were incubated with 50μM Mirin (MCE-HY-117693) for 4h prior addition of gemcitabine and maintained alongside gemcitabine for 24h.

Immunofluorescence images were acquired on a 63X-immersion oil lens equipped Zeiss microscope and Zeiss Blue software. Cells with ≥5 to 10 nuclear foci depending on the target protein were scored using CellProfiler (version 2.2.0, Broad Institute). At least three biological replicates (vehicle-, gemcitabine- and olaparib-treated) were analyzed.

For tumor tissues, 6μm cryosections were prepared from OCT embedded deep frozen tissue and mounted on Fisherbrand™ Superfrost™ Plus Microscope Slides (Fisher Scientific, Illkirch-Graffenstaden, France) and stored at -80°C until used. Tumor cryosections were immersed in 70% ethanol containing 0.1% SBB (Sigma Aldrich, Saint Quentin Fallavier, France) for 20min at RT to reduce autofluorescence and washed 3X 5min in PBS with 0.02% Tween 20. Subsequent immunostaining steps were identical to those applied on cell lines.

### Cell cycle determination and replication stress quantification by flow cytometry

7×10^5^ cells were grown for 24h in 10cm Petri dishes and subjected to a 30min pulse with 10μM EdU Click-it (ThermoFisher Scientific, les Ulis, France) prior gemcitabine treatment for 24h. After drug removal fresh medium was added and cell aliquots were collected at time points 0h, 8h, 24h, 48h. Cells were trypsinized and spun at 1200 rpm for 5min. Cell pellets were resuspended in the pre-extraction solution (PBS + 0.4% Triton X100) for 5min, then pelleted, fixed with 4%PFA in PBS for 30min at RT and permeabilized in PBS + 1% BSA + 1X saponin for 20min at RT. Cells were sequentially incubated for with the primary and the secondary antibodies for 60 and 30min at RT. Antibodies were diluted in PBS + 1%BSA + 1X saponin. Between primary and secondary antibody incubation cells were washed in PBS+1%BSA. After secondary antibody incubation cells were resuspended in PBS + 1%BSA + 1μg/ml DAPI for DNA counterstaining. FACS analysis was performed on a Gallios Flow (Beckman Coulter, Villepinte, France). Debris and doublets were excluded. Settings were identical in each channel and 10 000 cells were analyzed per aliquot. For the quantification of Sub-G1 fraction applied settings partially excluded debris and doublets. Quantification of apoptosis was done on cell pellets rinsed once in cold PBS and a second time in cold BBA buffer (10mM HEPES pH7.4, 110mM NaCl, 2.5mM CaCl2). Cells were resuspended in 50μl BBA buffer + 1μl AnnexinV (Sigma Aldrich, Saint Quentin Fallavier, France) and 0.02μg Propidium Iodide (PI) and incubated at RT for 30min. Prior cell analysis 200 μl PBS were added.

### Cell viability test

Experiments were performed in 96 well flat bottom plates (Starstedt, Marnay, France) with 1500 cells seeded in each well. After 24h 10μl serial drug dilutions were added in row of 8 wells. This was repeated in triplicates. Cells were exposed to the drug for 24h, gently pelleted and resuspended in fresh medium and left for approximately 2 cell cycle periods (48-56h), before addition of 10μl CCK8 (Cell Counting Kit-8, Tebubio Perray-en-Yvelines, France). Multi-well plates were incubated for 4h at 37°C, solutions in the wells gently homogenized and 450nM OD were measured on a Pherastar photometer (BMG Labtech Champigny-sur-Marne, France). Cell viability (%) = [(As-Ab) / (Ac-Ab)] × 100; As represents the absorbance of the experimental well, Ab the absorbance of a control well containing medium and CCK8, Ac the absorbance of a control well containing cells, medium and CCK8.

### Protein extraction and Western blotting

Protein extracts were prepared by lysing either tumor tissue or cell line pellets on ice for 30min in 50mM Tris-HCl pH7.4, NaCl 100mM, NaF 50mM, β-glycerophosphate 40mM, EDTA 5mM, Triton X100 1%, Aprotinin 10mg/ml, PMSF 100mM, Leupeptin 1mM, Pepstatin 1mM, followed by a short centrifugation to pellet debris. Protein concentrations were measured using the BCA kit (Fisher Scientific, Illkirch-Graffenstaden, France) SDS-PAGE gel electrophoresis was performed on 60µg protein samples subsequently transferred onto 0.22 µm nitrocellulose membranes (Amersham, Velizy-Villacoublay, France) and incubated overnight at 4°C with the primary antibody, after blocking for 1h in 10 % non-fat milk in TBST buffer (20mM Tris-HCl pH7.4, 150mM NaCl, 0.05% Tween20).

Antibodies used are listed in a separate section. Membranes were then washed and incubated with the appropriate secondary antibody in 5% non-fat dry milk in PBST for 2h at RT and revealed in Chemiluminescent HRP Substrate (Sigma Aldrich, Saint Quentin Fallavier, France).

### Genomic DNA double strand break determination by Pulse Field Gel Electrophoresis (PFGE)

Pellets of 1×10^6^ treated cells were incorporated in 1% agarose at 50°C and cast into molds (Biorad, Grabels, France) avoiding air bubbles. Samples in agarose plugs were incubated in lysis buffer (100mM EDTA pH8, 0.2% sodium lauryl sarcosine, 1mg/ml proteinase K) for 24h at 37°C, rinsed 3 times in wash buffer (20mM Tris, 50mM EDTA pH9) and positioned in the wells of a 1% agarose gel in 0.25X TBE. The gel was then run for 24h using pulsed electric field. DNA was labeled using Syber Green and revealed by UV. Fragmented DNA was quantified using the Image lab 6.0.1 software (Biorad, Grabels, France) and normalized relative to the non-fragmented DNA intensity.

### PDX models and in vivo treatment

TNBC PDX models establishment was as described (13). The study was reviewed and approved by the ethics committees for animal experimentations of the University of Montpellier (CEEA-LR-12028). PDX models were established from fresh tumor fragments obtained from the Pathology Department at the Comprehensive Cancer Center of Montpellier (ICM) after informed consent of the patients. Establishment of PDX models was reviewed and approved by the institutional review board. Approximately 50 mm^3^ PDX fragments were grafted subcutaneously into the flank of 3-4week old Swiss-nude female mice (Charles Rivers, Saint-Germain-sur-l’Arbresle, France). The present study comprised two experimental arms; vehicle and gemcitabine (Lilly, Fegersheim, France) comprising 8 mice per arm. When median tumor volume reached 100-150mm3, mice were randomly distributed in the two arms and treatment was started. Gemcitabine was injected intra-peritoneally (IP) twice/week for 4 weeks at 50 mg/kg. At treatment end, mice were euthanized, tumor samples collected for further histological analyses.

### Used antibodies

immunofluorescence/FACS : Mouse Anti-BrdU, BD Bioscience 347580 1/200, mouse anti-BRCA1, Santa Cruz sc-6954 1/300, rabbit anti-BRCA2, Bethyl A303-434A, 1/500, rabbit anti-phospho histone 3, CST 9701S, 1/50, rabbit anti-FANCD2, Abcam ab8928, 1/500, rabbit anti 53BP1, Novus Biological, NB100-34, 1/2000, rabbit anti RAD51, Merck PC130, 1/300, mouse anti-RPA32, Abcam ab2175, 1/200, rabbit anti-γH2AX, CST 9718S, 1/1000. Secondary antibodies; anti-rabbit alexa fluor 555, Abcam ab150075, 1/1000, anti-mouse alexa fluor 488, Abcam ab150113, 1/1000. Western blotting: rabbit anti-BRCA1, CST 9010? 1/500, rabbit anti-BRCA2, Bethyl A303-434A, 1/1000, rabbit anti-RAD51, CST 8875, 1/1000. Secondary antibodies; anti-rabbit HRP, CST 7076, 1/10000, anti-mouse HRP, CST 70745, 1/10000.

## Results

### BRCA1-deficient cell models exhibit increased sensitivity to gemcitabine

We determined the sensitivity to gemcitabine of 5 cell lines (4 TNBC and 1 ovarian cancer) with different *BRCA1* statuses; *BRCA1* wild type (BT-549 and SUM-159B1), *BRCA1*-mutated (MDA-MB-436) and full (OVCAR8) or partial (HCC38) *BRCA1*-promoter hypermethylation (14). In addition, we engineered by CRISPR-Cas9 a *BRCA1*-KO clone (SUM-159B1KO) from the SUM-159B1WT cell line (15). Noticeably, BRCA1-deficient cell lines MDA-MB-436, OVCAR8 and SUM-159B1KO showed distinctly lower IC50 to gemcitabine, compared with the BRCA1-proficient BT-549 and SUM-159B1WT and the partially *BRCA1*-hypermethylated TNBC cell line HCC38 (Figure 1A). Western Blotting analysis revealed that while BRCA1 protein expression was clearly detected in SUM-159, BT-549 and HCC38, it was absent in the *BRCA1*-deficient SUM-159B1KO, MDA-MB-436 and OVCAR8 cell lines (Supplementary Fig 1A). We also verified the protein expression levels of the important HR co-factors of BRCA1, BRCA2 and RAD51. Noticeably, all cell lines expressed similar BRCA2 and RAD51 levels. These data, thus, suggested that the superior sensitivity to gemcitabine could be linked to the absence of BRCA1 protein expression. Next, we focused our study on the isogenic SUM-159B1WT and SUM-159B1KO cell line pair and noted that the mortality induced by gemcitabine in SUM-159B1KO was five-fold higher than that of SUM-159B1WT cells (Figure 1B).

**Figure 1:**
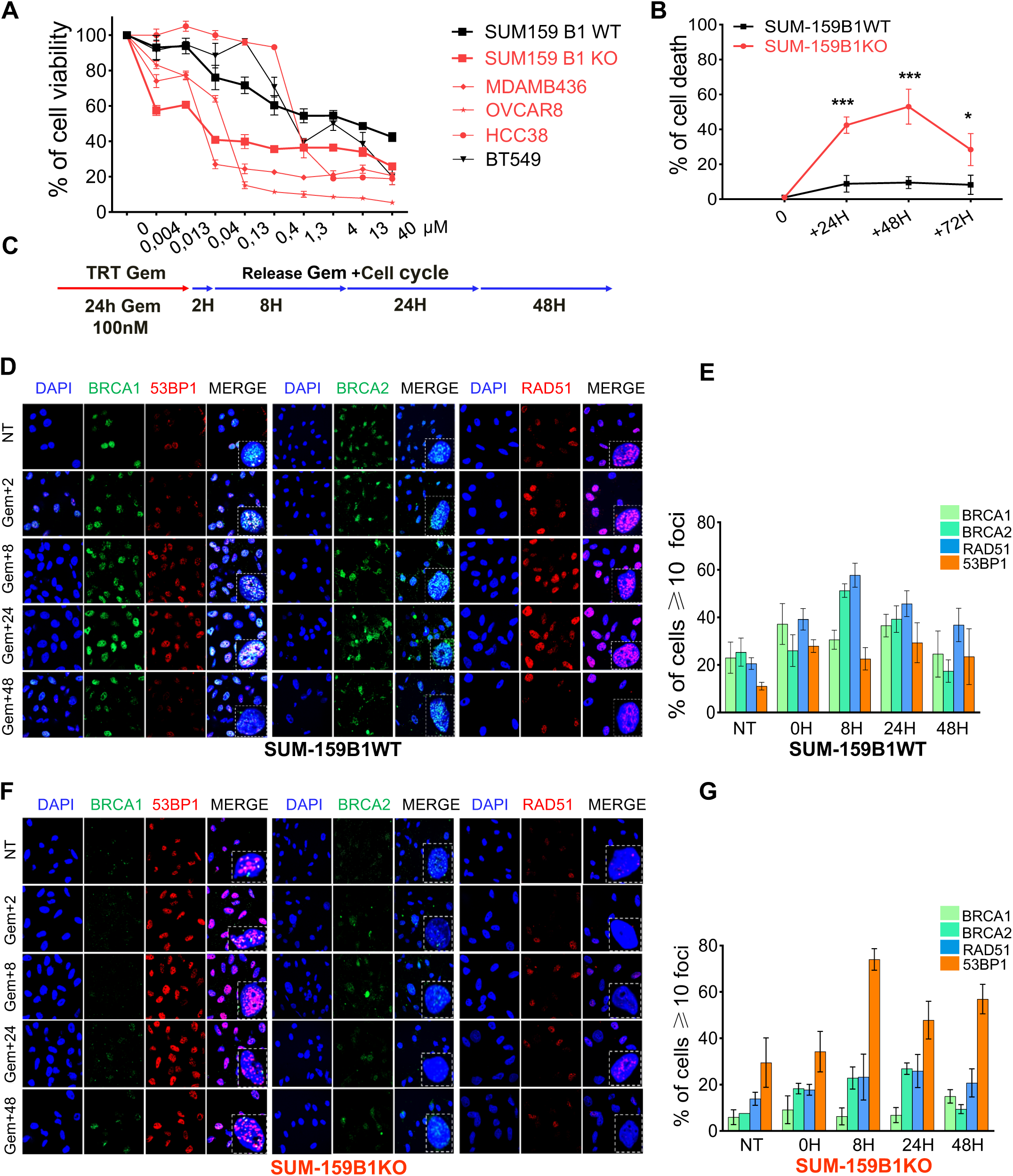
BRCA1-deficiency is associated with accrued gemcitabine mortality and absence of HR response. **A:** cell viability in response to increasing concentrations of gemcitabine, BRCA1-proficient cell lines are shown in black curves and BRCA1-deficient in red. **B:** cell mortality of gemcitabine treated SUM159B1WT (black) and SUM159B1KO (red) cells. Cell death was assessed by FACS quantification and combines AnnexinV+ and propidium iodide+ cells. **C:** treatment protocol applied here; cells were exposed to the drug for 24h, then left to recover in fresh medium for 48h before mortality was measured. **D, E**: immunofluorescence analysis and quantification of BRCA1, BRCA2, RAD51 and 53BP1 nuclear foci formation in SUM-159B1WT cells upon gemcitabine treatment. **F, G:** same as D, E but with SUM-159B1KO cells. Cells presenting more than 10 nuclear foci by immunofluorescence were scored positive and at least 300 cells were scored per slide. Boxes with dotted lines in the merge images correspond to magnification of typical nuclei. Statistical significance was assessed using the non-parametric Student t-test. P-values were considered as significant when *, ≤ 0.05 and highly significant when **, ≤ 0.01; ***, ≤ 0.005.

Thereafter, we applied a gemcitabine treatment protocol similar to that used for IC50 determination, *i.e.* 24h drug exposure followed by drug removal and analysis at different time points (Figure 1C). Cell cycle analysis of SUM-159B1WT and SUM-159B1KO cells treated with 100nM gemcitabine (IC50 of SUM-159B1WT) revealed that SUM-159B1WT cells reentered cell cycle 48h after removing the drug, while SUM-159B1KO further accumulated in S and G2/M cell cycle phases (Supplementary Fig 1B-C).

These data indicated that BRCA1-deficient cells treated with the replication poison gemcitabine undergo severe mortality combined with a strong G2/M blockage.

### BRCA1-deficient cells exhibit strongly reduced RAD51 and BRCA2 foci formation upon gemcitabine treatment

RAD51 and BRCA2, play important roles in DSB repair by HR, but have also been shown to be involved in the protection and repair of stalled replication forks (16). We, thus, performed an immunofluorescence time-course monitoring of nuclear RAD51 and BRCA2 foci formation levels, in conjunction with that of BRCA1 in gemcitabine-treated SUM-159B1KO and SUM-159B1WT. Furthermore, we scored foci numbers of 53BP1, a protein strongly binding to DSB sites that is actively removed and antagonized by BRCA1 (17). There is, thus, an inverted balance between BRCA1 and 53BP1 on DSBs and we were interested to assess this in situations of replication stress. Interestingly, whereas SUM-159B1WT treated with 100nM gemcitabine showed strongly increased BRCA1, BRCA2 and RAD51 foci formation, with a peak at 8h and a slow recess after 24h, in SUM-159B1KO BRCA1, BRCA2 and RAD51 foci formation was impaired. By contrast, 53BP1 foci were strongly increased (Figure 1F-G). We made very similar observations in the *BRCA1* mutated UWB1.289 ovarian carcinoma cell line, in comparison with its UWB1.289B1 variant ectopically expressing a WT *BRCA1* construct and observed impaired BRCA2 and RAD51 associated with increased 53BP1 foci formation in parental UWB1.289 cells (Supplementary Fig 1D-E).

### Gemcitabine-treated BRCA1-deficient models undergo persistent replicative stress

Cell populations subjected to replication stress exhibit an increase of γH2AX-positive and RPA-positive cell numbers, but in case of severe replication stress γH2AX-positive cells outnumber RPA-positive cells (18). We performed a FACS time course analysis of γH2AX-positive and RPA-positive cells in gemcitabine-treated SUM-159B1WT and SUM-159B1KO. We noted a temporary imbalance of γH2AX-positive and RPA-positive cell numbers at 0h and 8h in SUM-159B1WT (Figure 2A), whereas SUM-159B1KO exhibited a strong and persisting γH2AX/RPA imbalance (Figure 2B). The excess of γH2AX-positive relative to RPA-positive cells in gemcitabine-treated SUM-159B1KO in comparison with SUM-159B1WT was also detectable by immunofluorescence (Figure 2C-F). While the number of γH2AX-positive cells remained stable between 24h and 48h, the fraction of RPA-positive cells steadily decreased at 24h and 48h in SUM-159B1KO (Figure 2F). In identical conditions, SUM-159B1WT showed no imbalance between γH2AX-positive and RPA-positive cells, with decreasing numbers of γH2AX-positive and/or RPA-positive cells at 48H (Figure 2E). Similar observations were made in UWB1.289 and UWB1.289B1 cells, where *BRCA1*-deficient parental UWB1.289 cells showed a strong imbalance between γH2AX-positive and RPA-positive cells, whereas BRCA1-proficient UWB1.289B1 did not (Supplementary Fig 2A-D). These data, thus, supported that BRCA1-deficiency was associated with a severe and persistent replication stress upon gemcitabine treatment.

**Figure 2:**
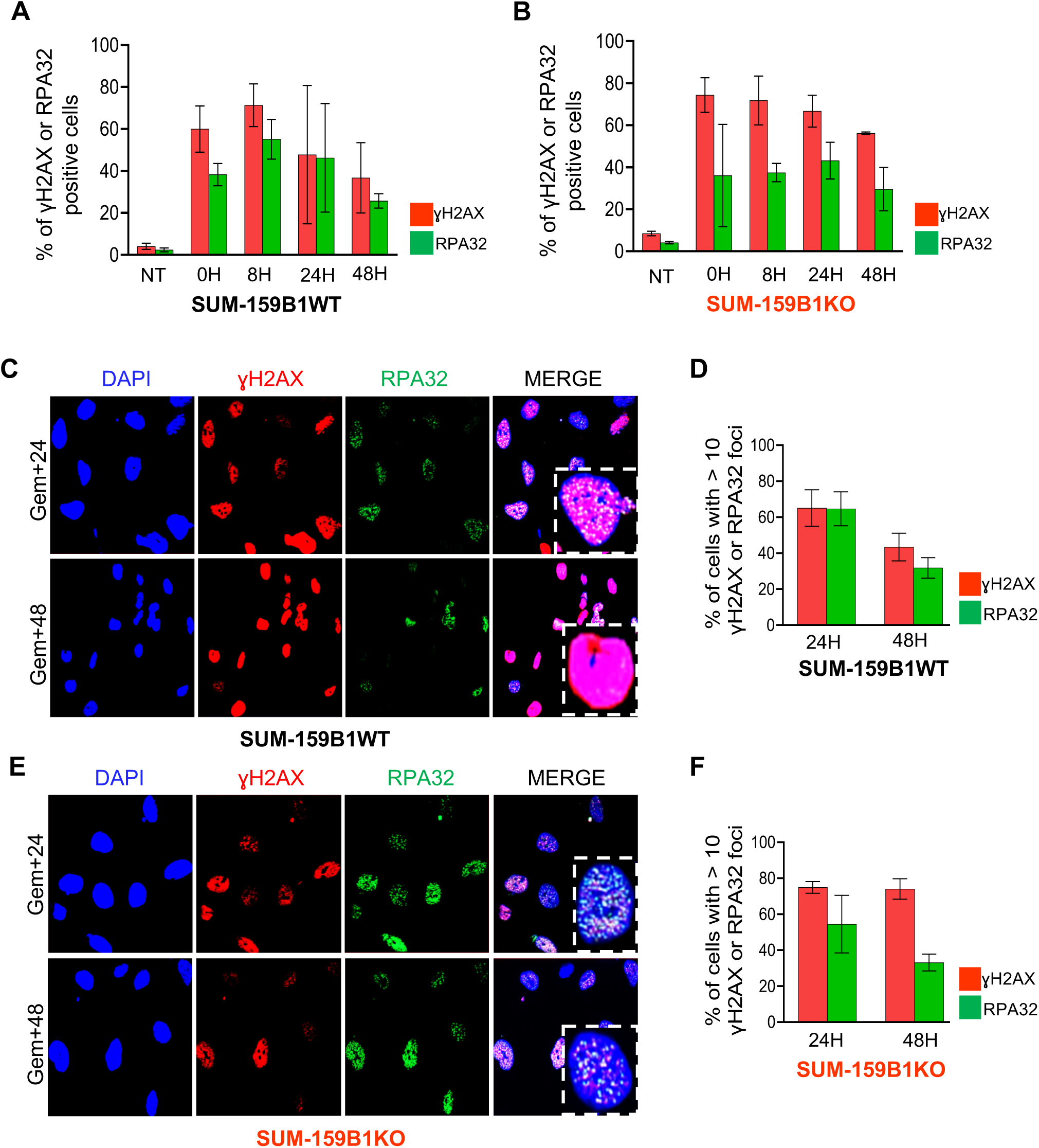
Gemcitabine treatment induces severe and persitent replication stress in BRCA1-deficient cell models. **A, B:** FACS analysis quantification of γH2AX-positive and RPA-positive cell fractions in SUM-159B1 (BRCA1-proficient) (A) and SUM-159B1KO (BRCA1-deficient) (B), note the imbalance between γH2AX- and RPA-positive cell numbers. **C, D:** SUM-159B1 showed no γH2AX/RPA imbalance in immunofluorescence (IF) analyses. Selected immunofluorescence sections of SUM-159B1WT stained with anti-γH2AX (red) and RPA (green) antibodies (C) and quantification of γH2AX-positive and RPA-positive cells by IF (D). **E, F:** SUM-159B1KO showed strong γH2AX/RPA imbalance by IF. Selected immunofluorescence sections stained with anti-γH2AX (red) and RPA (green) antibodies (E) and quantification of γH2AX-positive and RPA-positive cells by IF (F). At least 300 cells were scored per slide. Boxes with dotted lines in the merge images correspond to magnification of typical nuclei.

### BRCA1-deficient cells accumulate large quantities of ssDNA and pan-nuclear γH2AX-staining in absence of RPA signal

As revealed by BrdU immunostaining in non-denaturing condition, gemcitabine-treated cells accumulated ssDNA, with some cells showing intense BrdU staining (Figure 3A-C). The number of cells showing this intense BrdU staining was distinctly higher in SUM-159B1KO compared with SUM-159B1WT and, strikingly, most of these cells presented no signal of the ssDNA binding protein RPA signals. BrdU-positive/RPA-negative cell numbers gradually increased over time in SUM-159B1KO, reaching up to 65% of the cell population at 48h (Figure 3C, D). By contrast, in SUM-159B1WT BrdU-positive/RPA-negative cells did not exceed 20% (Figure 3B). Noticeably, cells with intense BrdU-staining were also characterized by a strong pan-nuclear γH2AX-staining (Figure 3E, F, Supplementary Fig3A). Furthermore, cells showing pan-nuclear γH2AX-staining presented no RPA foci, while cells with dotted γH2AX foci were RPA foci positive (γH2AX-positive/RPA-positive) (Figure 3G-I). Strikingly, the BrdU (ssDNA)/ pan-nuclear γH2AX-positive cells were mainly observed in SUM-159B1KO and other BRCA1-deficient cell models where their numbers strongly increased over time, reaching up to 70% of the BrdU-positive cells 48h after drug removal (Figure 3F, Supplementary Fig 3B-D).

**Figure 3:**
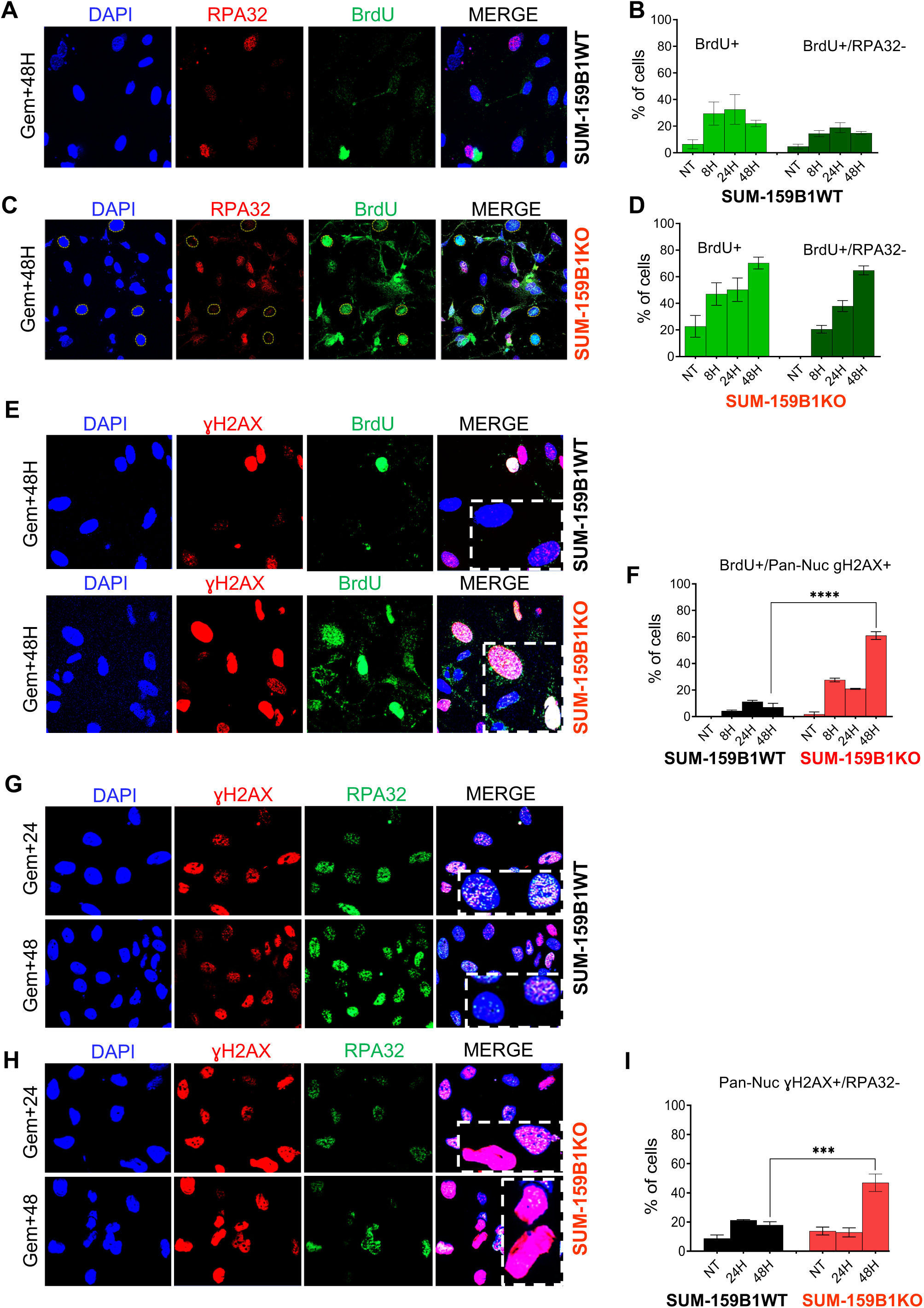
cells undergoing replication catastrophe are characterized by intense non-denaturing BrdU labelling, absence of RPA signal and pan-nuclear γH2AX staining. **A, B:** examples and quantification of RPA and BrdU immunofluorescence staining in SUM-159B1WT and SUM-159B1KO at different time lapses post gemcitabine removal. Yellow circles in B indicate cells showing no RPA signal and intense BrdU staining **C, D**: quantification of BrdU and RPA32 staining patterns in SUM-159B1WT (C) and SUM-159B1KO (D). **E**: examples of BrdU-positive and pan-nuclear γH2AX staining cells in SUM-159B1WT and SUM-159B1KO 48H after drug removal. **F**: quantification of the fraction of pan-nuclear γH2AX staining cells showing no RPA32 signal in SUM-159B1WT and SUM-159B1KO. **G, H**: examples of γH2AX and RPA co-staining in SUM-159B1WT and SUM-159B1KO. **I:** quantification of pan-nuclear γH2AX-positive and RPA-negative cells. Boxes with dotted lines in the merge images correspond to magnification of typical nuclei. Statistical significance was assessed using the non parametric Student t-test. P-values were considered significant when *, ≤ 0.05 and highly significant **, ≤ 0.01; ***, ≤ 0.005.

Remarkably, a large majority of SUM-159B1KO cells with pan-nuclear γH2AX staining, in addition of being devoid of RPA foci, were also RAD51 foci negative (pan-nuclear γH2AX-positive/RAD51-negative). By contrast, up to 80% of pan-nuclear γH2AX-positive SUM-159B1KO cells showed elevated numbers of 53BP1 and FANCD2 foci, in comparison with SUM-159B1WT where 53BP1 and FANCD2 positive cells did not exceed 30% (Figure 4A-F).

**Figure 4:**
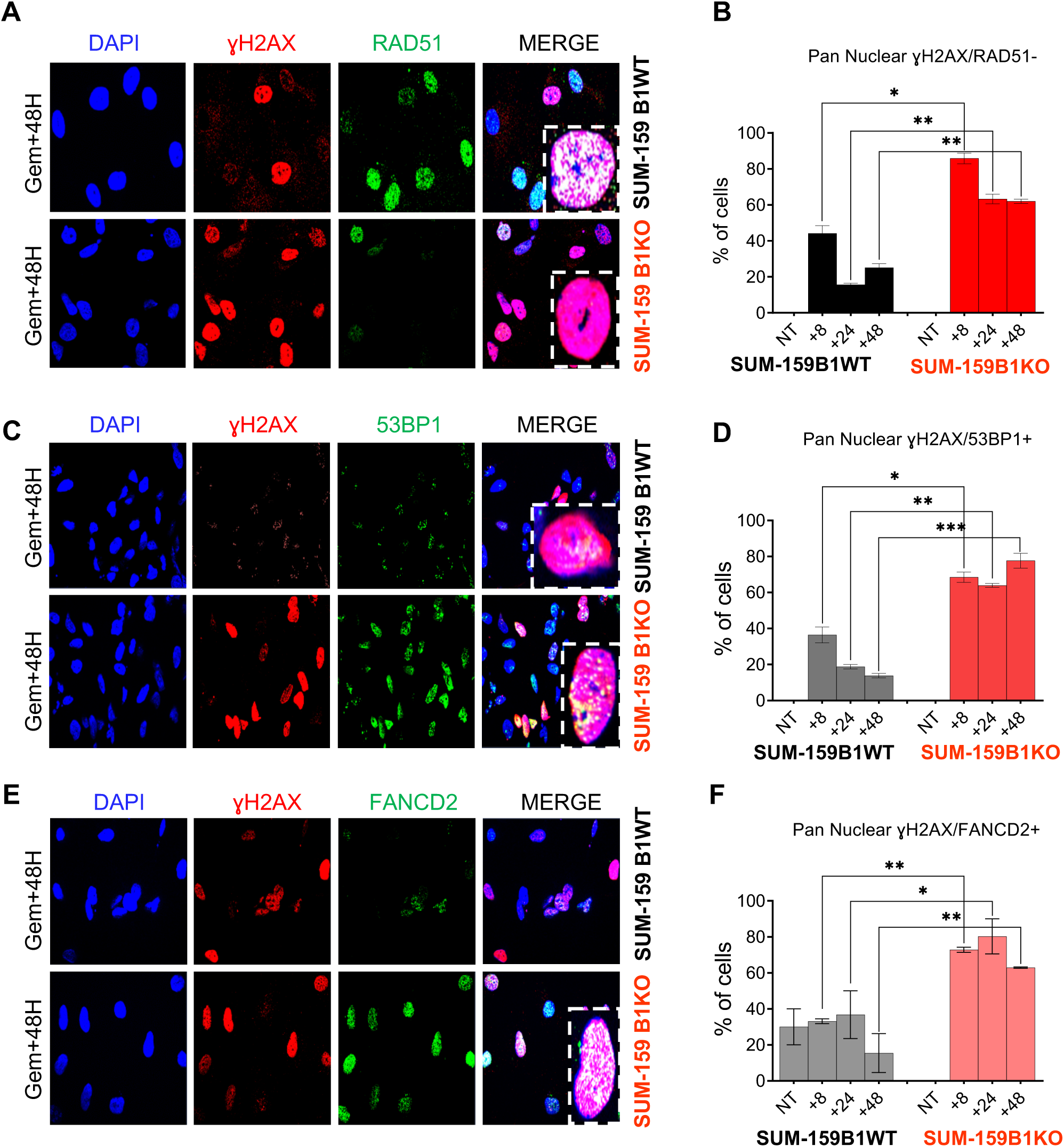
SUM159B1KO cells with pan-nuclear γH2AX staining show no RAD51 foci, but strong 53BP1 and FANCD2 signals. **A:** examples of γH2AX and RAD51 immunofluorescence staining 48H after drug removal. **B:** quantification of pan-nuclear γH2AX+/RAD51-cells in SUM-159B1WT and SUM-159B1KO. **C:** examples of γH2AX and 53BP1 immunostaining. **D:** quantification of pan-nuclear γH2AX+/53BP1+ cells in SUM-159B1WT and SUM-159B1KO. **E:** examples of γH2AX and FANCD2 immunostaining. **F:** quantification of pan-nuclear γH2AX+/FANCD2+ cells in SUM-159B1WT and SUM-159B1KO. Quantification was performed on at least 300 cells per section for each antibody. Boxes with dotted lines in the merge images correspond to magnification of typical nuclei. Statistical significance was assessed using the non-parametric Student t-test. P-values were considered as significant when *, ≤ 0.05 and highly significant when **, ≤ 0.01; ***, ≤ 0.005.

Our results thus indicate that, in BRCA1-deficient models, gemcitabine treatment resulted in a gradually increasing and persistent ssDNA accumulation devoid of RPA and/or RAD51 binding.

### MRE11 inhibition strongly reduced ssDNA accumulation and DSB levels in gemcitabine-treated SUM-159B1KO cells

The absence of RPA, RAD51 and reduced BRCA2 levels in gemcitabine-treated SUM-159B1KO cells constituted a context of poor fork protection favorable for uncontrolled MRE11 DNA resection (16),(19), (3). Consistent with this scenario treatment of SUM-159B1KO cells with 100nM gemcitabine + 50μM of the MRE11-inhibitor mirin resulted in a strong reduction of BrdU-positive and γH2AX-positive cells (Figure 5D-F). By contrast, mirin addition only induced marginal changes in SUM-159B1WT (Figure 5A-C).

**Figure 5:**
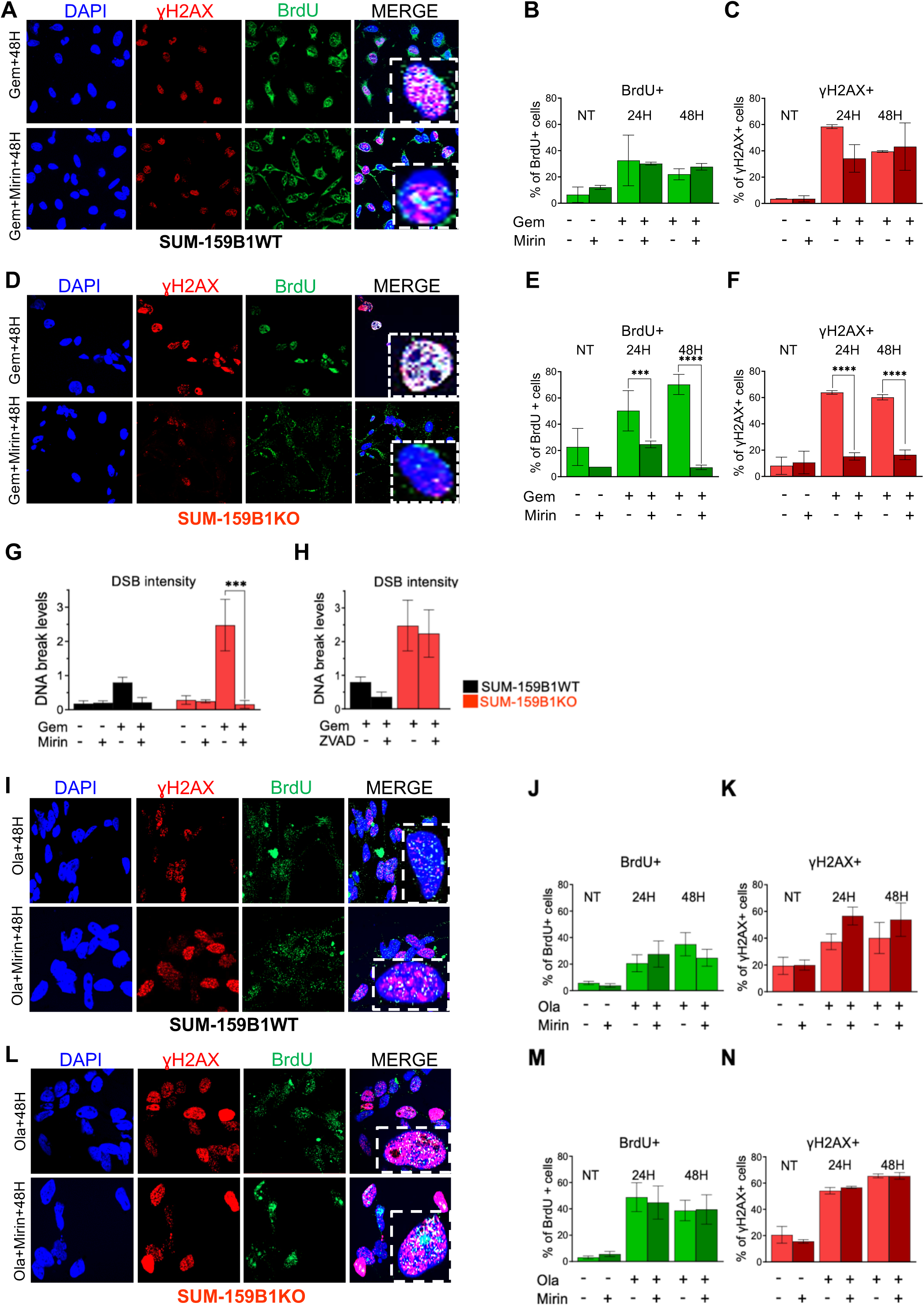
MRE11 is instrumental for ssDNA accumulation and onset of replication catastrophe in gemcitabine treated BRCA1-deficient cells. **A, B, C:** impact of mirin treatment on the number of BrdU and γH2AX positive cells in gemcitabine treated SUM-159B1WT. γH2AX and BrdU immunofluorescence staining in cells treated for 24h with Gemcitabine (top row) and 100nM Gemcitabine + 50μM mirin (bottom row) (A). Quantification of BrdU+ cells in the different conditions (B). Quantification of γH2AX+ cells (C). **D, E, F:** impact of mirin treatment on the number of BrdU and γH2AX positive cells in gemcitabine treated SUM-159B1KO. γH2AX and BrdU immunofluorescence staining in cells treated with Gemcitabine (top row) and Gemcitabine + mirin (bottom row) (D). Quantification of BrdU+ cells in the different conditions (E). Quantification of γH2AX+ cells (F). **G:** quantification of DSB breaks in SUM-159B1WT (black bars) and SUM-159B1KO (red bars) treated with gemcitabine +/-50μM mirin. **H:** quantification of DSB breaks in SUM-159B1WT and SUM-159B1KO treated with gemcitabine +/- 50μM Z-VAD. **I, J, K**: impact of mirin treatment on the number of BrdU and γH2AX positive cells in olaparib treated SUM-159B1WT cells. γH2AX and BrdU immunofluorescence staining in cells treated for 24h with 100 μM olaparib (top row) and 100 μM olaparib + 50μM mirin (bottom row) (I). Quantification of BrdU+ cells in the different conditions (J). Quantification of γH2AX+ cells (K). **L, M, N**: same as I, J, K in SUM-159B1KO cells. Boxes with dotted lines in the merge images correspond to magnification of typical nuclei. Statistical significance was assessed using the non-parametric Student t-test. P-values were considered as significant when *, ≤ 0.05 and highly significant when **, ≤ 0.01; ***, ≤ 0.005; ****, ≤ 0.001.

Furthermore, as unresolved replication fork stalling and breakdown can result in double strand DNA breaks (DSB), we determined DSB levels in gemcitabine-treated SUM-159B1WT and SUM-159B1KO by pulse field gel electrophoresis (PFGE). Gemcitabine-treated SUM-159B1KO accumulated at least two times more DSBs, in comparison with SUM-159B1WT in identical conditions. Furthermore, mirin addition strongly reduced DSB levels in SUM-159B1KO, while its effect was not as pervasive in SUM-159WT (Figure 5G), in line with the impact of MRE11 inhibition on ssDNA accumulation (Figure 5B, 5E). However, it has been proposed that, in BRCA2-defective models treated with olaparib, DNA breaks could result from apoptotic DNA fragmentation, rather than from replication fork breakdown (20). Hence, we verified the impact of 50μM ZVAD-FMK, a pan-caspase inhibitor, in gemcitabine-treated SUM-159B1KO and SUM-159B1WT. Noticeably, we observed no reduction in DSB levels, despite a strong reduction in apoptotic cell numbers (Figure 5H, Supplementary Fig 4A). Hence, our data indicated that uncontrolled MRE11 resection played a major role in the accumulation of ssDNA in gemcitabine-treated BRCA1-deficient models and was associated with severely increased DSB levels.

### While MRE11 resection plays a leading role in gemcitabine induced ssDNA accumulation, this is not the case in olaparib-treated cells

We further questioned whether MRE11 resection could also be seen in *BRCA1*-deficient cells upon olaparib treatment. Thus, the impact of MRE11 resection on ssDNA accumulation was tested in cells treated with 100μM olaparib (IC50 measured in SUM-159B1WT) +/-50μM mirin for 24h and, remarkably, both SUM-159B1WT and SUM-159B1KO showed no significant reduction of BrdU (ssDNA) nor γH2AX levels upon mirin addition. These data indicated that MRE11 resection did not play a major part in olaparib-induced replication stress (Figure 5I-N). This suggests that while gemcitabine treatment induces MRE11 hyper-resection dependent ssDNA accumulation and DSB, different mechanisms appear involved upon olaparib treatment.

### SUM-159B1KO cells undergoing replication catastrophe slip into mitosis resulting in mitotic catastrophe

*W*e had determined that gemcitabine-treated BRCA1-deficient cells tended to accumulate in G2/M (Supplementary Fig 1C) and formed micronuclei (MN), suggesting possible mitotic catastrophes.

Hence, we questioned whether gemcitabine-treated SUM-159B1KO cells had a tendency to slip into mitosis. To ascertain this, we synchronized the cells at the end of G1 by exposing them to 100nM of the CDK4/6 inhibitor palbociclib for 24h. Cells reached S phase synchronously 8h after palbociclib removal, time at which we added 1μM gemcitabine for 3h (Supplementary Fig 5A). After removing gemcitabine, FACS monitoring revealed that, while SUM-159B1WT resumed cell cycle at 96h, SUM-159B1KO accumulated in M phase at 72h and 96h (Supplementary Fig 5B, C). Interestingly, the cell cycle profile of gemcitabine-treated SUM-159B1KO was similar to that of cells treated with a combination of 50nM gemcitabine + 50nM of the CHK1 inhibitor PF-0477736 (Supplementary Fig 5D), suggesting that SUM-159B1KO presented a defective G2 to M checkpoint (21). Unscheduled progression into mitosis was in line with the large fraction of aberrant mitoses (multipolar divisions, ultrafine mitotic bridges or MN), representing up to 70% of the mitotic figures observed in gemcitabine-treated SUM-159B1KO cells (Figure 6A, B). In comparison, SUM-159B1WT presented only 20% aberrant mitoses (Figure 6B).

**Figure 6:**
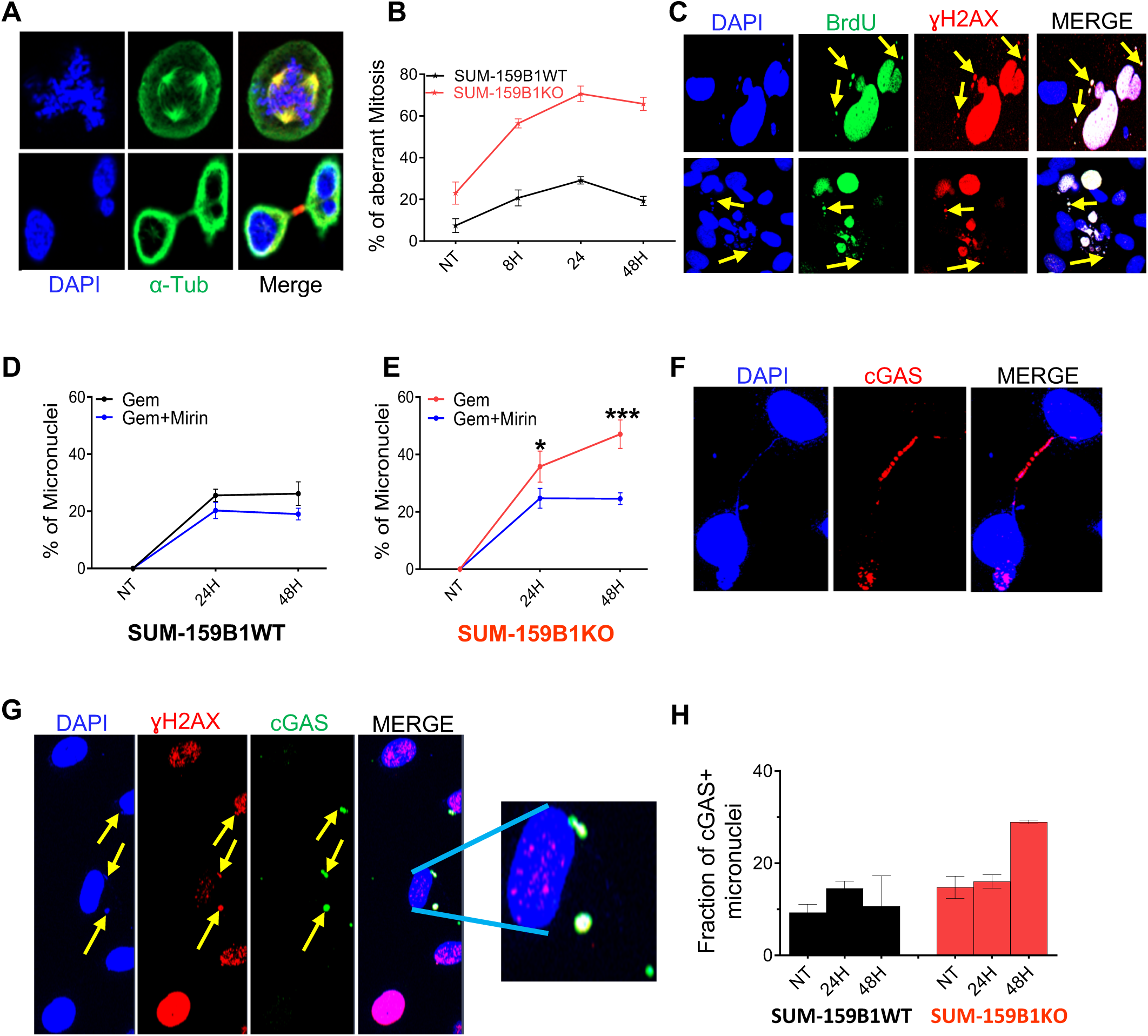
BRCA1-deficient cells undergoing replication catastrophe produce aberrant mitoses and cGAS-positive micronuclei. **A:** examples of aberrant mitotic figures in synchronized SUM-159B1KO cells treated with gemcitabine. **B:** fraction of aberrant mitoses in gemcitabine treated cells. Red SUM-159B1KO, black SUM-159B1WT. **C:** in SUM-159KO micronuclei are BrdU and γH2AX-positive. **D, E:** impact of 50μM mirin treatment on micronuclei numbers in gemcitabine-treated SUM-159B1 and SUM-159B1KO respectively. **F:** mitotic DNA bridges stain positive for cGAS in gemcitabine-treated SUM-159B1KO **G:** γH2AX+ micronuclei show cGAS staining in gemcitabine-treated SUM-159B1KO. **H:** quantification cGAS-positive micronuclei in SUM-159B1WT and SUM-159B1KO. Statistical significance was assessed using the non-parametric Student t-test. P-values were considered as significant when *, ≤ 0.05 and highly significant when **, ≤ 0.01; ***, ≤ 0.005.

### Micronuclei in SUM-159B1KO are linked to replicative catastrophe and sensed by cGAS

MN formation is linked to lagging mitotic chromosomes due to non-resolved genomic damage (22). Remarkably, we noted that MN in SUM-159B1KO presented both pan-nuclear γH2AX and strong non-denaturing BrdU staining, indicating they encompassed long stretches of ssDNA (Figure 6C). Due to the observed impact of MRE11 inhibition on ssDNA accumulation in SUM-159B1KO, we tested whether mirin treatment affected MN formation as well. Mirin treatment reduced MN numbers by 53% in SUM-159B1KO, while showing limited effect in SUM-159B1WT (Figure 6D, 6E).

It has been reported that mitotic bridges and MN could activate the cyclic GMP-AMP synthase (cGAS) sensing response, due to cytoplasmic exposure of DNA (22),(23). Accordingly, we noted that, in gemcitabine-treated SUM159B1KO, ultrafine DNA bridges and up to 30% BrdU-positive micronuclei showed a clear cGAS signal (Figure 6F, 6G). By contrast, the cGAS signal was significantly less prevalent in the few MN detected in SUM159B1WT (Figure 6H).

Our data, thus, indicate that BRCA1-deficient cells undergoing replication catastrophe slip into mitosis and produce MN associated with a strong cGAS sensing.

### Gemcitabine responsive TNBC PDX models show increased γH2AX and cGAS staining

The strong increase in γH2AX levels assorted to pan-nuclear staining observed in cell models undergoing massive ssDNA accumulation led us to verify whether γH2AX staining levels could be related to gemcitabine sensitivity in breast cancer patient derived xenograft (PDX) models. To this mean, we selected four TNBC PDX models whose sensitivity to two injections per week of 50mg/kg gemcitabine for four weeks was determined (Figure 7A-7D). One model (15b0018) progressed, two (b3804, b1995) were stabilized, while the last one (b4122) clearly regressed within the four weeks of treatment. The day after the last drug injection residual tumors were sampled and immunolabeled for γH2AX and cGAS expression. γH2AX and cGAS positive cells were quantified in gemcitabine and mock treated tumor sections and their numbers were highest in PDX b4122, which responded best to gemcitabine and lowest in PDX 15b0018 that progressed under treatment. Noticeably, PDX b3804 and b1995, whose growth was stabilized under gemcitabine treatment, showed intermediate γH2AX and distinctly lower cGAS staining levels (Figure 7E-7H). These *in vivo* data are, thus, consonant with our cell culture model observations and suggest that elevated sensitivity to gemcitabine treatment could be marked by increased γH2AX and cGAS staining.

**Figure 7:**
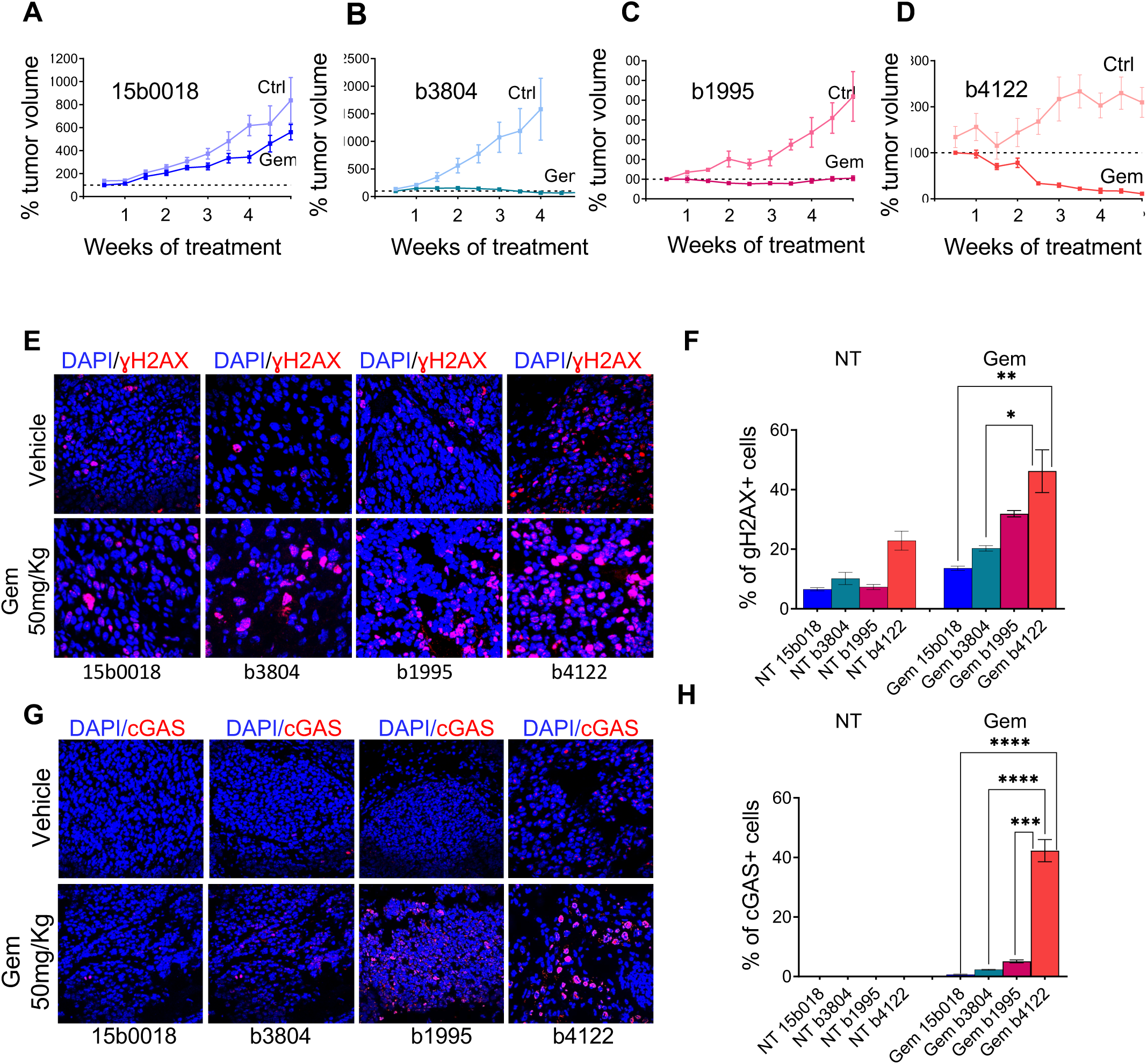
TNBC PDX showing good response to gemcitabine exhibit increased γH2AX and cGAS staining. **A, B, C, D:** growth curves of PDX b15b0018 , b3804, b1995 and b4122; black lines mock, red lines gemcitabine-treated. About 50mm3 of tumors were grafted subcutaneously to 8 mice in each experimental arm and treatment started when tumor volume had reached 150mm3 on average. Two injections/week of 50mg/kg gemcitabine were administered by IP injection for 4 weeks. Mice in the control arms were injected with the vehicule. **E, F:** γH2AX immunostaining and quantification of positive cells**. G, H:** cGAS immunostaining and quantification of positive cells. Statistical significance was assessed using the non-parametric Student t-test. P-values were considered as significant when *, ≤ 0.05 and highly significant when **, ≤ 0.01; ***, ≤ 0.005.

## Discussion

Here we show that *BRCA1*-deficient cell models are particularly sensitive to the replication poison gemcitabine in comparison with their BRCA1-proficient counterparts. Upon gemcitabine treatment *BRCA1*-deficient models encounter massive ssDNA accumulation and elevated cell death. Because these cells showed strong and persistent imbalance of γH2AX- and RPA-positive cells, this process was clearly reminiscent of replication catastrophe, an acute and lethal form of replication stress, originally described in association with ATR impairment (18). Replication catastrophe was proposed to result from an excess of ssDNA accumulation, which titrates available RPA in the cell and results in severely impaired protection of stalled replication forks (18). By contrast, *BRCA1*-proficient models only showed transient γH2AX/RPA imbalance and modest mortality. Furthermore, we report, for the first time to our knowledge, that gemcitabine-treated *BRCA1*-deficient cells massively accumulate ssDNA in absence of concomitant RPA and/or RAD51 signals. Remarkably, the number of cells exhibiting this phenotype increased over time, reaching up to 70% of the cells 48h after drug removal. These results suggested that ssDNA levels produced in gemcitabine-treated BRCA1-deficient models exceeded those required for RPA titration and, thus, involved further events. The fact that addition of the MRE11 inhibitor mirin resulted in a strong reduction of ssDNA levels accumulated in gemcitabine-treated SUM-159B1KO cells clearly indicated the leading role of uncontrolled resection by MRE11. MRE11 resection of nascent strands has been proposed to be associated with replication fork reversal (11), (24). Replication fork reversal, a response to acute replication fork stalling that puts replication forks on hold in the perspective of repair, is a remodeling process that creates structures potentially vulnerable to nucleolytic attack (25). Because of impaired RAD51 recruitment, reversed forks appear particularly exposed in *BRCA*-deficient cells (16), (11), (26), (12) (27). Our data showed that neither RPA nor RAD51 were detectable, thus, constituting a context clearly favorable to uncontrolled resection. We further noted that, while no RAD51 and RPA signals were detected, 53BP1 and FANCD2 foci numbers were increased in gemcitabine-treated *BRCA1*-deficient cells, with no apparent effect on fork protection. It is, however, of note that MRE11 is insufficiently processive to support resection of long DNA stretches, like those observed in gemcitabine-treated BRCA1-deficient models. It has been shown that this requires the involvement of further nucleases such as EXO1 and/or WRN/DNA2 (27), (28). MRE11 recruitment to nascent DNA stretches thus appears to act as a priming event in strand resection. Moreover, the recent finding, that MRE11 binds to gemcitabine incorporated into the nascent DNA strand to actively remove it, constitutes an interesting lead, possibly explaining the imported role of MRE11 observed in gemcitabine-treated *BRCA1*-deficient models (29). By contrast, we noted that 24h treatment with 100μM olaparib did not result in mirin sensitive ssDNA accumulation in SUM-159B1KO cells. Our results are in agreement with previous work showing that DNA synthesis repriming is preferentially activated and fork reversal repressed upon olaparib or PARPi treatment (20), (30). DNA repriming downstream of stalled forks, by way of the primase directed DNA polymerase PRIMPOL, leaves ssDNA gaps and MRE11 has no documented role in this process (31) (3).

Remarkably, *BRCA1*-deficient cells undergoing massive ssDNA accumulation showed intense pan-nuclear γH2AX staining, instead of discrete γH2AX foci. Moreover, the numbers of cells showing this phenotype increased brutally between 24 and 48h after drug removal. By contrast, BrdU-positive cells numbers increased regularly between 8 and 48h, suggesting that pan-nuclear γH2AX staining could represent late, possibly final, stages of replication catastrophe. The shift from γH2AX foci to pan-nuclear staining could illustrate massive DNA breakage resulting from replication fork breakdown and correspond, as proposed by Moeglin and coauthors (32), to a pre-apoptotic signal. Our data showing that addition of Z-VAD to gemcitabine-treated SUM-159B1KO strongly reduced pan-nuclear γH2AX staining cell numbers support the link with apoptosis (Supplementary Fig 4B).

This led us to question whether the level of γH2AX staining in gemcitabine-treated PDX TNBC models could reflect their level of sensitivity to the drug. Our analysis indicated that γH2AX levels were indeed increased, both in terms of positive cell numbers and staining intensity, in PDX models showing better response to gemcitabine treatment. These results suggest that the intensity and pattern of γH2AX staining in post-treatment patient biopsies could be informative of the responsiveness of a tumor to replication poisons. However, this will need to be verified on larger cohorts of PDX models and patient biopsies to be ascertained.

Somewhat to our surprise, *BRCA1*-deficient TNBC cells suffering from replication catastrophe, despite robust ssDNA accumulation, reached M phase and produced aberrant mitoses, particularly, micronuclei (MN) showing intense BrdU labeling. It is of note that MRE11 inhibition reduced BrdU-positive MN numbers. Hence, it is most likely that BrdU-positive MN originated from lagging chromosomes bearing long stretches of ssDNA (33). The presence of fine chromatin bridges in the aberrant mitotic fields of gemcitabine-treated SUM159-B1KO cells supported this hypothesis. In addition, we noted that BrdU and γH2AX labeled MN also presented a strong signal of the cytoplasmic DNA sensor cGAS. Cyclic GMP–AMP synthase (cGAS) is a component of the innate immune system (34), which is frequently activated in tumor cells and was shown to react with MN (23). It has been proposed that cGAS sensing of DNA contained in MN could be linked to mitotic bridge sensing prior MN formation and nuclear envelope reconstitution (22). It must be noted that cGAS sensing of ssDNA loaded MN appears somewhat heterodoxical, as cGAS has been shown to specifically bind double strand DNA (dsDNA), not ssDNA. However, the presence of interstitial dsDNA stretches cannot be ruled out and could thus explain cGAS staining observed on mitotic bridges and MNs. cGAS DNA sensing is an early signal of the interferon inflammatory immune response, which in cancer could trigger an antitumor immune response, especially as cGAMP has been shown to participate to paracrine spread and activation of the tumor microenvironment (22). It is of note that the PDX model responding best to gemcitabine treatment also presented a clear cGAS staining.

Hence, our data suggest that gemcitabine treatment of BRCA-deficient TNBC could be beneficial to patients because of; (1) its impact on tumor cell survival, (2) the possible induction of a secondary antitumor immune response.

### Conflict of interest

None of the authors has a conflict of interest to disclose

### Author contribution

Imene Tabet; Conceptualization, investigation, validation, methodology, drafting and editing the paper. Esin Orhan; investigation, validation, methodology, editing the draft. Ermes Candiello; Conceptualization, investigation, editing the draft. Carolina Velazquez; investigation, data curation, validation, methodology. Lise Fenou; resources, validation, data curation, methodology. Beatrice Orsetti; methodology, resources. Geneviève Rodier; methodology, validation, editing the draft. William Jacot; resources, editing the draft. Cyril Ribeyre; investigation, data curation, validation, methodology, editing the draft. Claude Sardet; funding acquisition, supervision, editing the draft. Charles Theillet; Conceptualization, Supervision and coordination, writing and editing the paper, funding acquisition,

### Ethics approval

Patients whose tumor was used to generate PDX signed informed consent and the preclinical assay was reviewed and approved by the ethics committees for animal experimentations of the University of Montpellier (CEEA-LR-12028) and received the approval number 25612.

### Data availability

The data presented in this study are available on reasonable request from the corresponding author.

## Acknowledgements

The authors sincerely acknowledge Dr Philippe Pasero (IGH) and Dr Isabelle Jariel (IRCM) for their fruitful discussions during the course of this work and constructive comments on the manuscript. Furthermore, sincere thanks are due to the staffs of the animal facility at IRCM for their constant support and expert help.

## Grants and financial support

This work benefited from the following financial support: Institutional support from the Institut National de la Santé et de la Recherche Médicale, Astra-Zeneca contract # 2018-02069, Institut National du Cancer PRTK-2017 MODUREPOIS, Ligue Nationale Contre le Cancer ‘Comité régional Occitanie-Est’ 2021-R22031FF and the SIRIC Montpellier Cancer Grant INCa-DGOS-Inserm_12553, LabMuse Epigenmed METABOHOX from the University of Montpellier.

## Author information

Ermes Candielo, present address; MBC-DBMSS Institute – IRCCS, Torino, Italy.

Carolina Velazquez, present address; Gynecological Oncology Laboratory, Department of Oncology, KU Leuven and Leuven Cancer Institute (LKI), 3000 Leuven, Belgium.

## Notes

### Competing Interest Statement

The authors have declared no competing interest.

### Summary of Updates

We have added data showing that, while gemcitabine treated BRCA1-deficient models undergo MRE11 dependent accumulation of ssDNA, this does not occur when the identical models are treated with the PARP-inhibitor olaparib. We also extended our in vivo PDX dataset and present data on four TNBC PDX models instead of two as originally shown. Moreover, the text and illustrations have been improved in order to enhance the clarity of the argument.

