## Supplementary material for "Replication poison treated BRCA1-deficient breast cancers are prone to MRE11 over-resection resulting in single strand DNA accumulation and mitotic catastrophe": Sup Figure 1-4

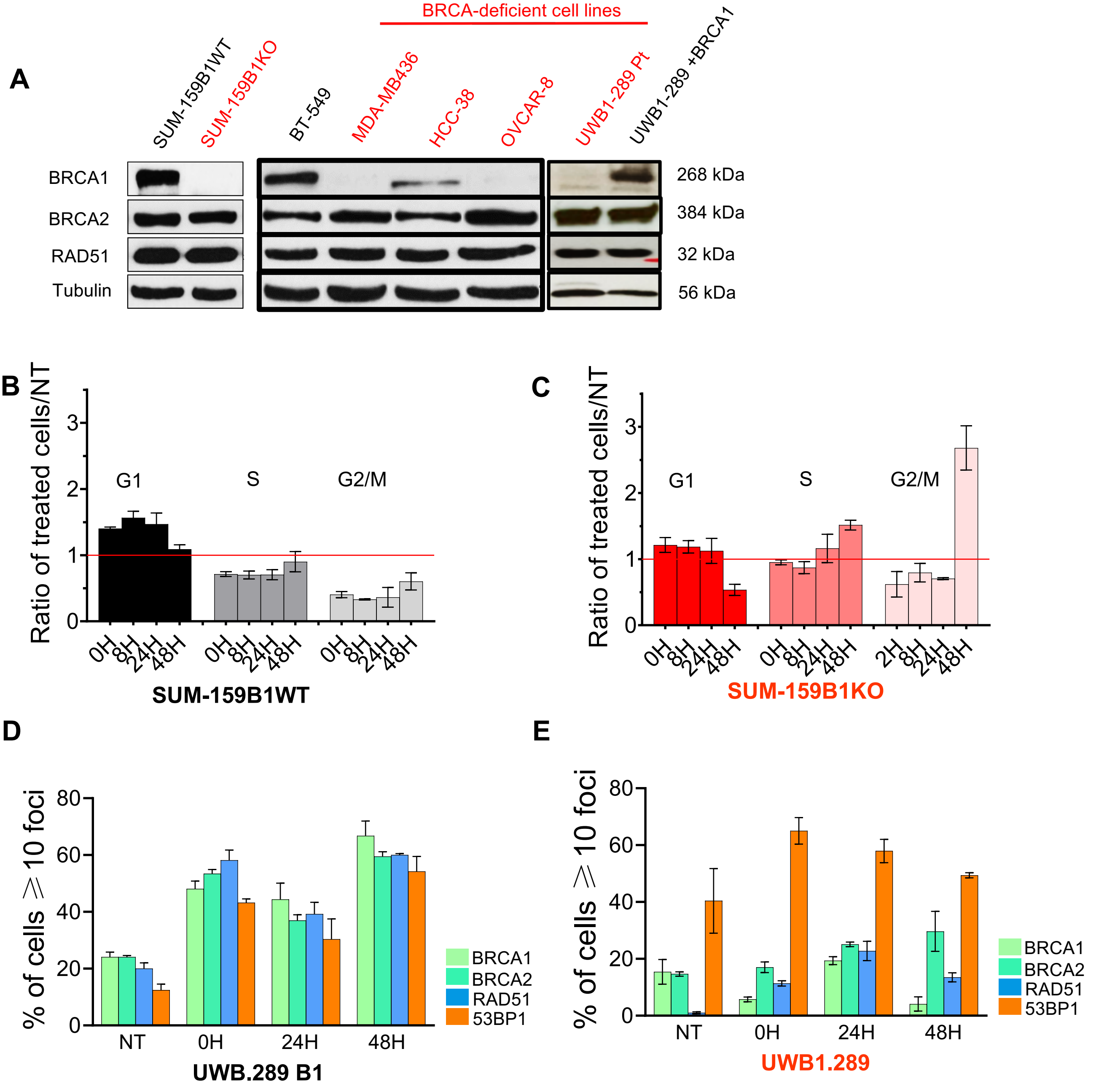


**Supplementary Fig 1**: **A**: western blot analysis of BRCA1, BRCA2 and RAD51 protein expression in the different TNBC cell lines used in this work. The BRCA1 status of the analyzed cell lines is indicated on top of the gel. **B, C**: cell cycle profile determination by FACS of gemcitabine treated SUM-159WT and SUM-159KO models; cells in respectively G1, S and G2/M phases were quantified immediately after (OH), 8H, 24H, 48H removal of the drug and normalized according to levels observed in untreated cells (red line). **D, E**: quantification of BRCA1, BRCA2, RAD51 and 53BP1 nuclear foci positive UWB1.289B1 (ectopic BRCA1 expression construct) and UWB1.289 (parental BRCA1-mutated cell line). Conditions and markers were quantified as in Figure 1C-F.

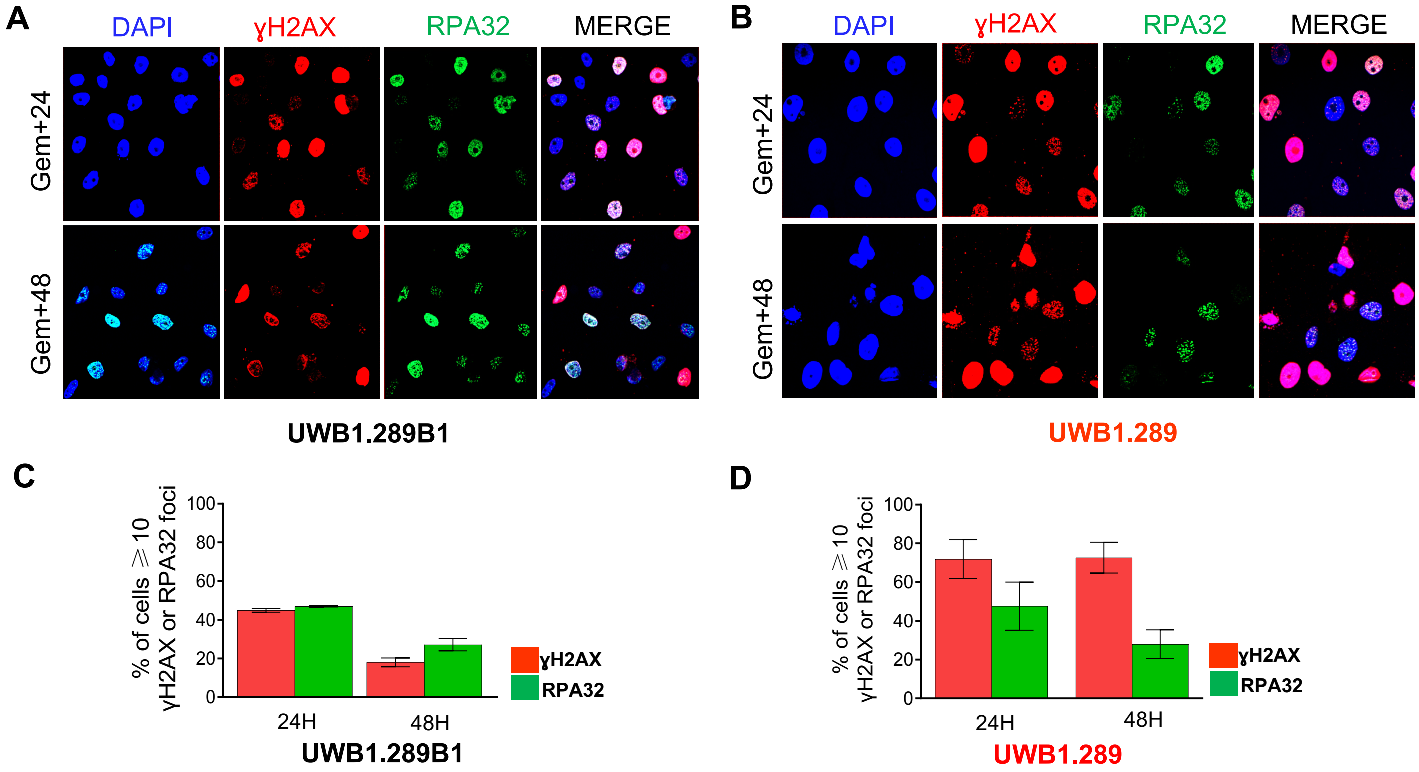


**Supplementary Fig 2**: **severe γH2AX/RPA imbalance in the BRCA1 mutated UWB1.289 in comparison with UWB1.289B1 (ectopic BRCA1wt expression). A, B**: selected sets of γH2AX (red) and RPA (green) immunofluorescence staining in UWB1.289 and UWB1.289B1 respectively. **C, D**: quantification of γH2AX and RPA32 positive cells in UWB1.289 and UWB1.289B1 immunofluorescence slides.

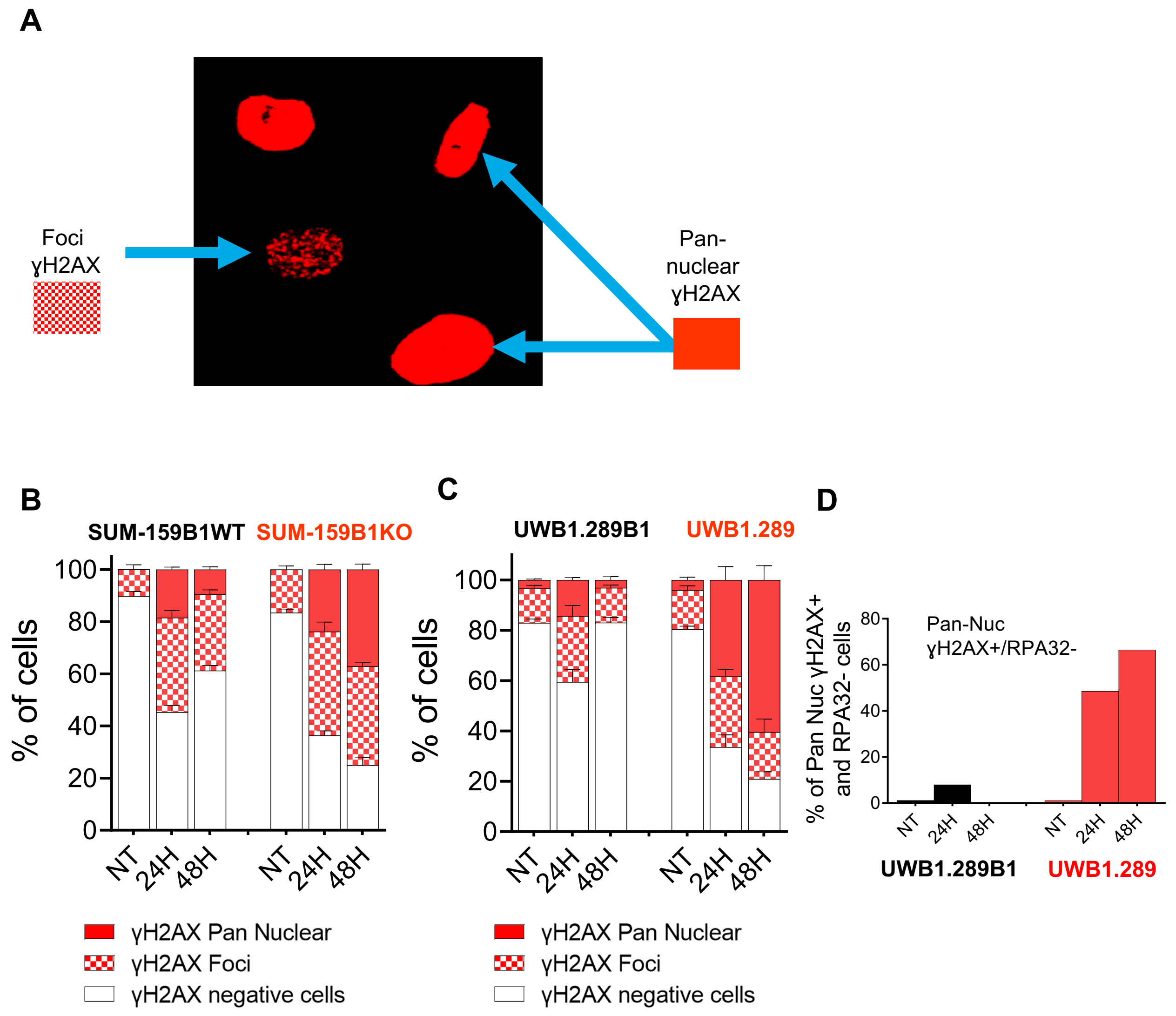


**Supplementary Fig 3: gemcitabine treated BRCA1-deficient cells accumulate increasing numbers of pan-nuclear γH2AX staining cells**. **A:** example of pan-nuclear vs. dotted γH2AX IF patterns. **B:** quantification and time course of γH2AX positive cells in SUM-159B1WT and SUM-159B1KO. **C:** quantification and time course of γH2AX positive cells in UWB1.289B1 and UWB1.289. **D:** quantification of pan-nuclear γH2AX+/ RPA32- cells in UWB1.289B1 and UWB1.289.

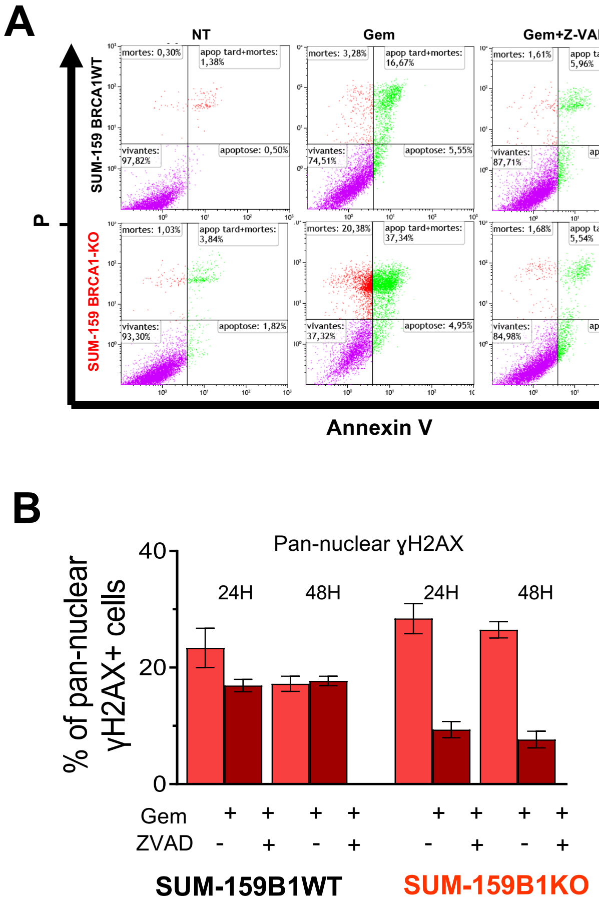


**Supplementary Fig 4: A:** Z-VAD-FMK treatment strongly reduces the fraction of annexin V positive in gemcitabine treated cells. **B**: impact of ZVAD treatment on pan-nuclear γH2AX staining cell numbers.

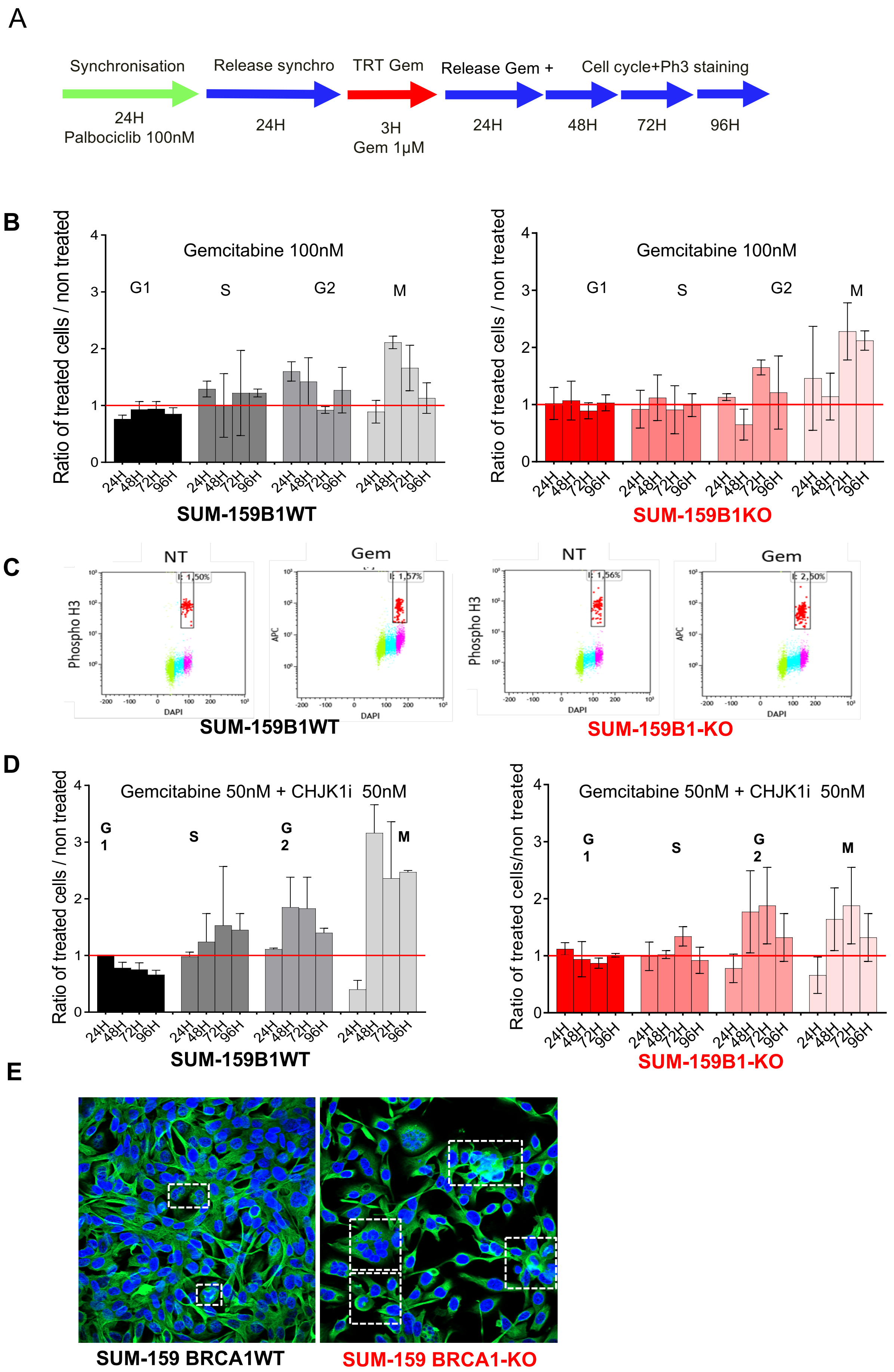


**Supplementary Fig 5: A**: cell synchronisation and treatment scheme. **B**: time course analysis of cell cycle patterns of gemcitabine-treated SUM-159B1WT and SUM-159B1KO cells. **C**: quantification of M phase cells (anti-Phospho-Histone3 positive) in gemcitabine-treated 24H after drug release **D**: cell cycle changes of synchronized SUM-159B1WT and SUM-159KO cells treated with a combination of 50nM gemcitabine + 50nM PF-0477736 (CHK1i). Cells slip into mitosis. **E:** SUM-159B1KO cells treated with gemcitabine produce aberrant mitotic figures.
